# Epigenetic transgenerational effects on RNAi in Arabidopsis

**DOI:** 10.1101/2020.04.05.026641

**Authors:** Qüan’an Hu, Jérôme Ailhas, Todd Blevins, Ulrich Klahre, Franck Vazquez, Michael Florian Mette, Frederick Meins

**Author notes:** The majority of works of this article were performed in the FMI, Switzerland.

## Abstract

Epigenetic states and even certain environmental responses in nematodes, mammals and seed plants can be inherited over successive generations. RNAi in plants is an epigenetic form of RNA degradation that is normally reset early in development. In contrast, competence, i.e., the susceptibility of a transgene for RNAi, is believed to be genetically determined and faithfully transmitted over many generations. Unexpectedly, we found that the same yellow fluorescent protein (YFP) transgene in a monogenic, homozygous line of *Arabidopsis* exhibited two epigenetic states of RNAi: type I, which is initiated in rosette leaves and is stable in later developmental stages, and, type II, which is initiated after floral induction and is unstable. The incidence of type I plants descended from type I parents decreased over three successive generations of self-fertilized plants. By the third generation, a substantial fraction of plants from type I parents exhibited either the type II phenotype or remained high-YFP expressing throughout sporophytic development, indicating that silencing had been completely lost. Similarly, plants descended from type II parents showed a progressive shift from the type II phenotype and a complete loss of silencing. These results show that the silencing types (or states) of parental plants have significant influence on both silencing determination and silencing types (or states) of their offspring, however, this influence fades through several generations of self; as regardless of the types of the parental lines, competence for RNA silencing tends to loss after generations. Thus, we propose that RNA silencing in plants can be the target of transgenerational effects.

While the underlying mechanisms are still unknown, our findings in the *Arabidopsis* RNA silencing system complement the current understanding of heritable RNAi in *C. elegans,* that this phenomenon is likely a common mechanism in Eukaryote systems.

## Introduction

Epigenetic states and even certain environmental responses in nematodes, mammals and seed plants can be inherited over successive generations[1–16]. These intriguing transgenerational effects have medical ramifications [17], potentially may be used in crop breeding[18], and could have important evolutionary consequences [11,19,20]. Some forms of epigenetic inheritance such as genomic imprinting, are normally reset in the next generation [21,22]. Other forms including paramutation [6], epimutation [23], some cell-heritable changes in growth factor requirement [2] and stress effects on somatic homologous recombination [3] persist with varying stability in successive generations.

RNA interference (RNAi), also called posttranscriptional gene silencing, is an evolutionarily conserved form of RNA silencing that results from the degradation of cognate mRNAs guided by 21-26-nucleotide-long small interfering RNAs (siRNAs) [24,25]. The genetic consequences of RNAi, from one generation to another, was firstly observed in *C. elegans;* that RNAi status can be heritable independent of the dsRNA trigger [7,8,10,11,26]. However, in plants, RNA triggered RNAi (RNA silencing, or post-transcriptional gene silencing) was not observed to be inheritable, but rather, RNA-triggered transcriptional gene silencing can be inherited independently of the RNA trigger and requires MET1 for maintenance [27]. RNAi of plant transgenes is usually a stochastic phenomenon that shows limited epigenetic inheritance [28]. Only some plants in a genetically homogeneous population show silencing and patterns of silencing vary from plant to plant. The site where RNAi is initiated combined with the subsequent local- and systemic-spread of silencing generate patterns that depend on the transgene insert, developmental stage of the plant, and environmental factors[29,30]. Once established, the silent state usually persists throughout sporophytic development, but is then reset to the non-silent state early in embryogenesis. Thus, in plants, RNAi of a stable transgene is a heritable, but potentially reversible epigenetic form of RNA degradation [31]. This has led to the current view in plants, that competence, i.e., the susceptibility of a transgene for RNAi, is determined by a genetic predisposition that is faithfully inherited, whereas the silent state is reset and then stochastically initiated in each generation. The present study challenges this view: We show that the same Arabidopsis transgene can exhibit multiple states of competence for RNAi; that different silent states of parental lines have an effect on the frequency of different silencing states of their offsprings, indicate RNA silencing is target for epigenetic transgenerational effects; plus that competence for RNAi can be progressively lost in successive generations.

## Materials and methods

### Plant material

Wild-type and transgenic Arabidopsis thaliana Col-0 plants were raised in soil under long-day conditions (16 h 10,000 lux light at 21° C /8h dark at 16° C) in growth chambers at 65-70% relative humidity. The homozygous GFP transgenic EG5 [32] shows transcriptional gene silencing (Blevins, unpublished). Transformants were generated by the floral dip method [33] with *Agrobacterium tumefaciens* strain GV3101 carrying the vector plasmid pAS6.

### Construction of the YFP-intron expression vector pAS6

pAS6 (Figure 1) carries the YFP coding region of pEYFP (Clontech) interrupted with the synthetic, 85 bp, plant intron Syn7 [34] regulated by a double-enhancer Cauliflower mosaic virus 35S RNA promoter (P35S2) and a nos terminator in the T-DNA binary vector pBI121 (Clontech, Switzerland). Syn7 was amplified with the primers 5′- GAGCTGAAAGggtaagattatcgatatttaaattatttatttcttcttttc-3′ and 5′-TGAAGTCGATGCctgcatgatatcataaaaccatgga-3′ specifying the splicing junctions and then inserted into the YFP coding region by overlapping PCR, in combination with these primers and the primers 5′-CTTGTACAGCTCGTCCATGCCGA-3′ and 5′- TGCCATGGTGAGCAAGGGCGA-3′, respectively.

**Fig. 1.**
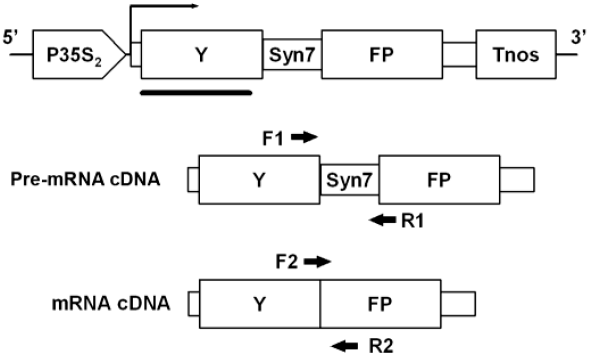
Maps of the *35S_2_-YFP_ivs_* expression casette of pAS and the expected spliced and unspliced cDNAs. The maps, which are not drawn to scale, show the positions of the 35S promoter (P35S_2_), the nos terminator (Tnos), and the YFP coding region interrupted by the intron *Syn7.* The horizontal bar indicates the hybridization probe used for RNA-blot analysis; the arrows indicate the approximate position of the primers used for quantitative RT-PCR of spliced and unspliced transcripts. We verified that the primer pair F1 and R1 amplified the 123 bp fragment expected for unspliced pre-mRNA cDNA and that the primer pair F2 and R2 amplified the 185 bp fragment expected for spliced mRNA cDNA (data not shown).

### RNA analyses

Total RNA was isolated from leaf tissue using Trizol reagent (Gibco BRL, Basel). Total RNA and small RNAs fractions were analyzed by RNA-blot hybridization as described by [35] using DNA and RNA probes for the full-length YFPivs sequence. Semi-quantitative real-time PCR was performed and the results normalized for expression of the At3g18780/ACTIN2 gene as described by [36]. cDNAs were synthesized from 500ng of total RNA as the template with gene specific-primers and M-MuLV Reverse Transcriptase (Fermentas, Germany). Quantification was performed in a iQcycler apparatus (Biorad, Germany) using SYBR green and the primers

F1: 5′-CTTCTTCAAGGACGACGGCAACTA-3′ and R1: 5′-CTTGAAGTCGATGCctgcatg-3′ for primary, unspliced RNA, and

F2: 5′-CCGCATCGAGCTGAAAGGCATC-3′ and R2: 5′-TTGATGCCGTTCTTCTGCTTGTCG-3′, for spliced mRNA (Fig. S1). Nuclear run-on transcription experiments were performed as described by (van Blokland et al. 1994) using probes for the coding regions of 35S2-YFPivs, ACTIN1 (At2g37620), and UBIQUITIN4 (At5g20620).

### DNA analysis

Total DNA was isolated from lower leaves with a DNeasy Plant Mini Kit (Qiagen, Germany). The genomic position of the T-DNA insert was determined by tail-PCR as described by (Liu et al. 1995) and BLAST searches of the TAIR 6.0 database. The structure of the T-DNA insert was analyzed by Southern blot hybridization of genomic DNA as described by [32] with a DIG-labelled DNA probe for a 730 bp region of the YFPivs coding region (Figure 1). Promoter methylation was analyzed by cleavage with methylation-sensitive restriction enzymes and Southern blot hybridization as described by [37] using the 32P-labeled sense RNA probe which covers the first 360 bp of YFP coding region.

## Results

### The same 35S2-YFPivs transgene exhibits two types of RNAi

To detect transgenerational effects on silencing, we examined the incidence and pattern of silencing in successive generations obtained by self-fertilization of Arabidopsis plants carrying a well-characterized transgene. The line we used, 23, is homozygous for the yellow fluorescent protein (YFP) reporter gene 35S2-YFPivs, which is interrupted by the synthetic intron Syn7, known to be very efficiently spliced [34] (Figure 1). Segregation test of Kanamycin resistant and YFP fluorescent indicate that this line has a singly transgene insertion. Southern-blot hybridization and TAIL-PCR suggest that line 23 contains 3 copies of tandem direct repeat inserted in chromosome 4 at position 9626285 (Figure S1a,b). Line 23 and most progeny obtained by self-fertilization (i.e., the S1 generation) exhibit yellowish and reddish fluorescence indicative of YFP silencing (Fig. 2a,b). RNA-blot hybridization confirmed a marked reduction of YFP mRNA in silent leaves relative to that of high-YFP expressing leaves (Figure 3a). Comparison of steady-state mRNA levels with those of nascent YFP RNA measured by nuclear run-on transcription (Figure 3b) showed that silencing of YFP expression in line 23 results from RNAi. This conclusion was confirmed by the finding that 21 nt long small interfering RNAs diagnostic for RNAi (Hamilton et al. 1999) accumulated in silent leaf tissues, but not in high-YFP expressing leaf tissues (Figure 3c).

**Fig. 2.**
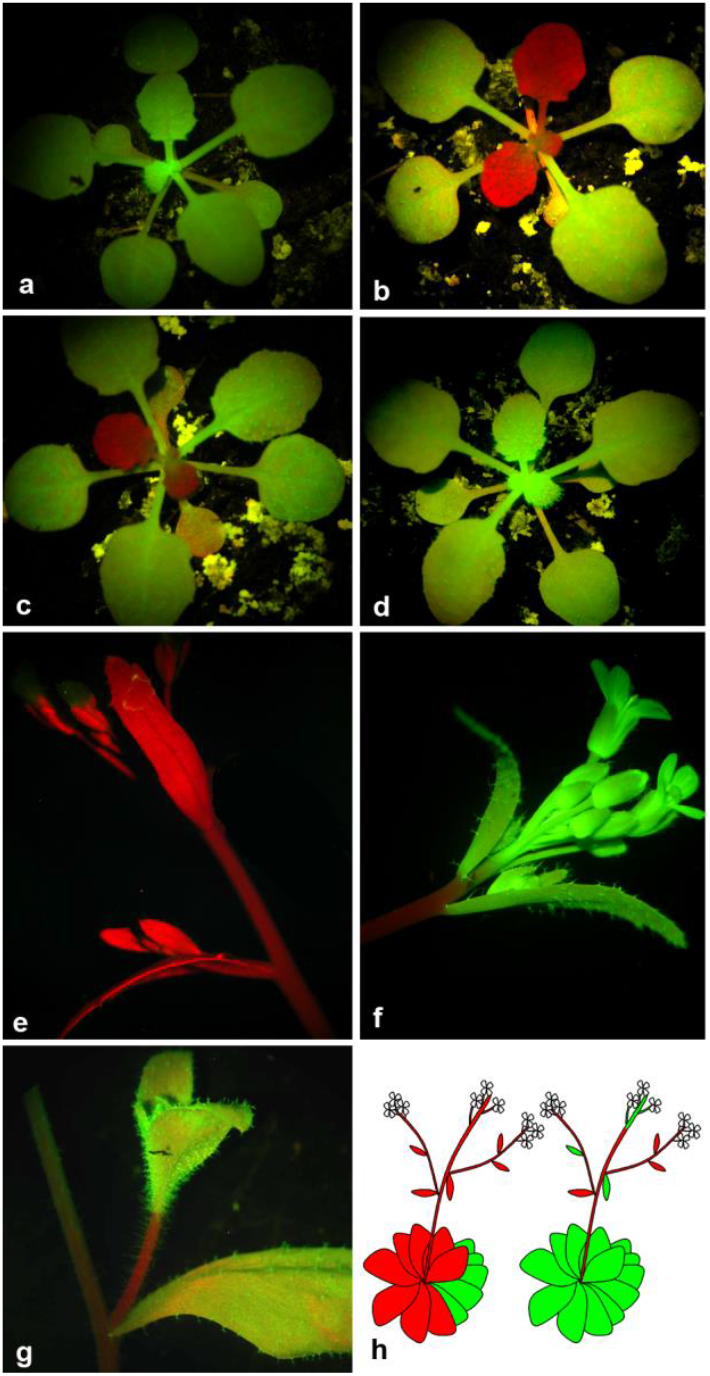
Representative images of line 23 plants showing type I and type II silencing. Rosette-stage high-YFP expressing plant (a), type I plants showing silencing starting in leaf 5 (b) and leaf 6 (c), and a type II plant at a comparable stage (d) that does not silence until late in development. Uniform silencing of the inflorescence of a type II plant (e). Unstable silencing with reversion in the inflorescence (f) and upper leaves (g) of type II plants. Summarized features of type I and type II silencing (h). Plants photographed under UV illumination show greenish-yellow fluorescence due to YFP or red chlorophyll autofluorescence in the absence of YFP. Bars: 5mm

**Fig. 3.**
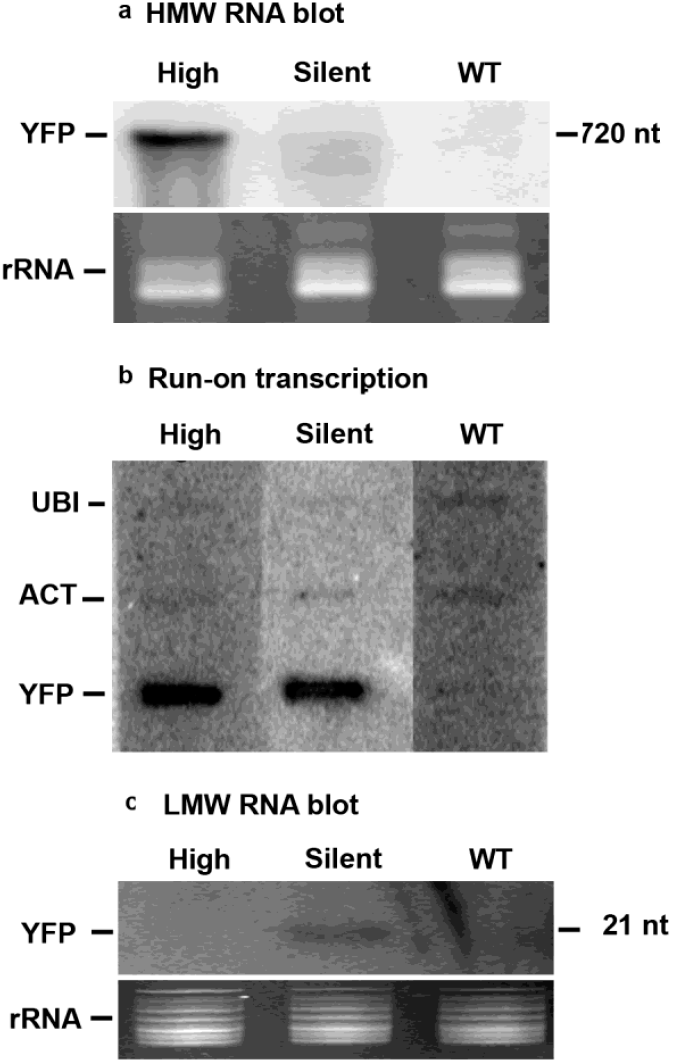
Evidence for RNAi of the *35S_2_-YFP_ivs_* transgene in line 23. Nuclei and RNA were prepared from high YFP expressing (high) and silent leaves of a line 23 plant, and from leaves of a wild-type (WT) plant. (a) RNA-blot hybridization of 10 μg of high-molecular-weight (HMW) RNA. (b) Slot-blot hybridization of nuclear run-on transcription products. (c) RNA-blot hybridization of 10 μg of low-molecular-weight (LMW) RNA. Blots were hybridized as indicated with probes for YFP_ivs_ (Fig. 1); *ACTIN1 (ACT1),* and *UBIQUITIN4 (UBI)* transcripts. The rRNA loading standards (b and c) were stained with ethidium bromide. Sizes in nucleotides (nt) indicated on the right were estimated from standards.

Figure 4 shows the onset of silencing in three self-fertilized generations (S1, S2 and S3) of known pedigree. Almost all, 91 of 93 S1 plants showed typical patterns of silencing starting in rosette leaves 3 to 13 and most frequently in leaf 5 or 6 (Fig. 2b,c). Although some leaves showed progressive, vein-dependent silencing indicative of local spreading [31], most emergent leaves were uniformly silent. Once initiated, silencing was stable and persisted in subsequent leaves, internodes, and inflorescences formed later in development (Fig. 2b,c,e). Besides, rosette leaves of two S1 plants did not show silencing (Fig. 2d); Silencing was first detected in internodes and cauline leaves after floral induction and was unstable, i.e., some cauline leaves and even entire inflorescences arising from silent nodes showed high levels of YFP (Fig. 2f,g). To summarize, in S1 generation plants, onset of RNAi stochastically distributed at different development stage. Depending on if or not the onset RNAi will persist to the whole plant, especially if or not to the flowers, We classify them to two types: an early onset in rosette leaves, before bolting will persist to the flowers, as stable form (type I); and, a later onset in internodes and cauline leaves, after bolting, could not persist to the flowers, as unstable form (type II) (Fig. 2h).

**Fig. 4.**
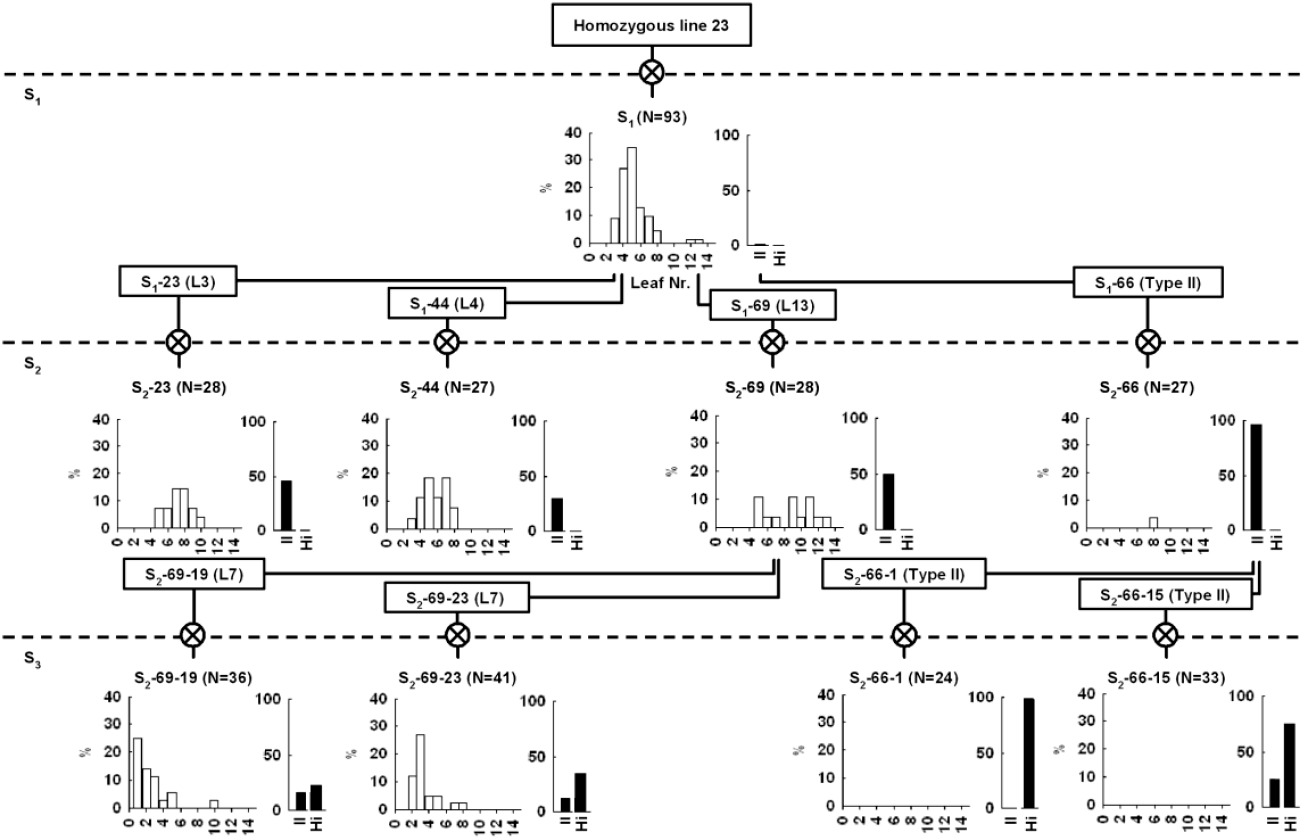
Incidence of RNAi in successive generations (S_1_-S_3_) obtained by self-fertilization of individual plants homozygous for the *35S_2_-YFPivs* transgene. Onset of silencing of type I plants (left graph) expressed as % of N siblings exhibiting silencing starting with the true leaf numbered 1-14. The incidence, expressed as % of N siblings, of type II plants showing unstable silencing (right graph, II) and high-YFP expressing plants (Hi). Plants were scored for YFP- and chlorophyll-fluorescence at flowering stage. Boxes show the generation, identifying number, and silencing type, or for type I plants, the first leaf showing silencing (L) of individual plants.

### Onset of silencing in type I plants is not heritable

We examined the inheritance of the stochastic patterns of RNAi in type I plants by comparing the onset of silencing in rosette leaves of progeny obtained from individual parents (Fig. 4). The onset of silencing in S2 progeny of parents showing early onset in leaf 3 (plant 23) and leaf 4 (plant 44) was similar to that of the parental S1 generation. In contrast, S2 progeny of plant 69 showing onset in leaf 13 gave a distribution skewed toward later onset of silencing. Nevertheless, plants in the same lineage, i.e., type I S3 progeny of S2 siblings 69-19 and 69-23, which showed onset of silencing in leaf 7 (plants), gave distributions similar to that of the S1 generation. Therefore, the silencing onset time in progeny was not consistently linked to that of the parent. Although predisposition for stochastic silencing of rosette leaves is heritable, the specific time of onset in parents is not generally inherited in the next generation.

### Parental epigenetic states have a genetic influence on that of their offsprings

We also examined the influence of parental silencing types to proportion of silencing types of their offspring, by testing progeny of selfed type I and type II plants (Fig. 4). If different silencing states are completely reset after meiosis, we should expect similar proportions of type I and type II plants in both groups of offspring. If silencing states is stably inherited, the proportion of type I plants should be maintained in lines selected for type I silencing and the proportion of type II plants should increase in lines selected for the type II silencing. As indicated in Fig.4, selection for type II silencing greatly increased the incidence of type II plants. In contrast, selection for type I silencing decreased the incidence of type I plants from 98% in the S1 generation to ca. 50% in the S2 and S3 generations. Moreover, selection for either silencing types frequently resulted in S3 plants showing a uniformly high YFP expression. Therefore, silencing types of parental plants have a genetic impact on silencing types of their offspring, plus a strong bias for transgenerational changes from type I to type II and the eventual loss of silencing.

Several lines of evidence suggest that these transgenerational effects reflect altered epigenetic inheritance of competence for RNAi: 1) The changes are heritable. Environmental effects are minimal because different generations assayed at the same time in independent experiments gave similar results. 2) The plants were homozygous progeny of one and the same transformant carrying a single transgene locus and, hence, were presumably genetically equivalent. 3) Conspicuous genetic alterations at the transgene locus are unlikely since the number and size of restriction fragments detected by Southern-blot analyses using probes for the T-DNA insert were the same for S1, S2, and S3 plants (Figure S1c). 4) The heritable changes are potentially reversible. Although infrequent, type II individuals can give rise to type I progeny (Fig. 3). 4) Finally, cotyledons and the first true leaves invariably exhibited the high YFP phenotype, confirming that RNAi is reset early in the next sexual generation [31]. This implies that competence for RNAi type rather than RNAi per se changes in successive generations. Moreover, the resulted offspring differences in between selfing type I and selfing type II indicate that parental epigenetic types have a genetic consequence to their offspring.

### Transgenerational changes in transcriptional activity

Transgenerational effects in both animals and plants often involve altered transcriptional activity and DNA methylation [5]. We compared transcriptional activity in S1 and S3 plants. As quantitative run-on nuclear transcription assays were not feasible with the limited amounts of tissues available, we used quantitative RT-PCR to measure levels of primary, unspliced YFPivs pre-mRNA and mature, spliced YFP mRNA. Our rational was that a previous study had shown that primary, unspliced transcripts in tobacco provide a good estimate of transcription rate that is not appreciably affected by RNAi [35]. Because this did not exclude the possibility that RNAi can impair pre-mRNA processing as reported for transgenic tomato and petunia [38,39], in the present study we assayed rosette leaves 1-4 that do not show RNAi.

Expression of unspliced transcripts in leaves of S1 type I plants was significantly higher (t-test of means) than in type I, type II and high-YFP expressing plants in the two lineages of S3 plants (Fig. 5a). Expression in the S3 type I plants was significantly higher than in either sibling type II plants or high expressing plants, but expression in S3 type II and high-expressing plants in the same lineage (S3-69) did not differ significantly. Expression of mature, spliced YFP mRNA was variable, but showed a positive linear correlation with the expression of unspliced transcripts (Fig. 5b, P < 0.01). If unspliced transcript levels in lower leaves truly reflect transcriptional activity, then our results suggest that a generation-dependent decrease in transcription could account for the shift from type I to type II competence, but not necessarily for differences between type II and high-expressing plants.

**Fig. 5.**
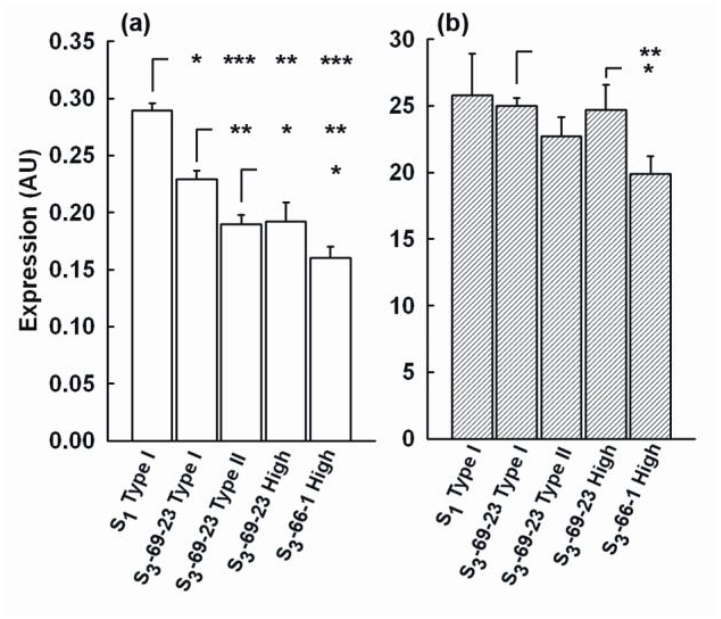
Transgenerational effects on transcriptional activity. Accumulation of unspliced primary YFP transcript (a) and mature, spliced YFP mRNA (b) in type I, type II and high-YFP expressing plants. Data are shown for three samples of RNA prepared from pools of high-YFP expressing lower rosette leaves (leaf 1 to 4), each from 15 sibling plants. RNA was measured by quantitative RT-PCR using primers for spliced and unspliced YFP transcripts. Values are normalized based on ACTIN2 mRNA and expressed in arbitrary units (AU). Error bars: ± SEM for three samples. Significance levels (t-test of means): *, P<0.05; **, P<0.01; ***, 0.001.

As DNA methylation of promoter regions is frequently associated with transcriptional repression in plants [5], we used Southern analysis of genomic DNA to detect methylation in CG, CNN, and CNG contexts at several restriction sites in the promoter of p35S_2_-YFPivs. No changes in methylation were detected in comparisons of type I and type II plants over three generations or between silent and high-expressing plants in the S3 generation (Figure S2a,b).

## Discussion

We have identified RNAi in Arabidopsis as a target for epigenetic transgenerational effects. RNAi in plants was triggered if the transcript level of a transgene surpassed a genespecific threshold[40]. The timing and spread of RNAi in individual plants is known to depend on several parameters including the plant species, the nature of the transgene, the structure of the transgene insertion, and environmental factors

In this experiment, using YFP as a marker, we recorded onset of silencing at different development stages of different individual plants, of the same homogenous transgenic line. We found that silencing may occur in earlier 3^rd^ leaf development stage, as well as in later development stage after bolting. We found in the specific transgenic line, that the earlier silencing in true leaves before bolting is “stable” and persist to all later on tissues, including flowers; while the later silencing after bolting may exhibit instability, that silencing may occur but could not persist to all later on tissues, including flowers. Depending on its “stability” and if or not persisting to the flowers, we classified silencing to “type I” and “type II”.

While onset of RNAi is stochastically, competence for RNAi is heritable, but can change from type I to type II, or be lost at variable rates in successive generations. Moreover, during such transition, silencing status of the parental lines have an effect on that of their offspring. Offspring of Type I silencing plants have higher chances or competences for onset of type I silencing. This provides an experimental example that epigenetic status, even though the regulation is at posttranscriptional level, have a significant effect on traits (silencing status in this case) of their offspring.

Genetic studies point to extensive cross-talk between transcriptional gene silencing (TGS) and RNAi [41–45]. Our measurements of unspliced transcripts in plants of known pedigree and silencing phenotype provide evidence that the rate of transcription of a transgene can decrease in successive generations and is higher in type I plants than in type II plants or high expressing plants. These findings and earlier studies showing that the same transgene in tobacco can exhibit RNAi and TGS [46,47], suggest that TGS and RNAi can contribute in varying degree to inactivation of the same transgene and that this is not a special feature of a particular transgene or plant species.

The molecular basis for the effects we observed is not known. Mourrain et al. (2007) have evidence that a complex nitrate reductase transgene locus of tobacco generates dsRNAs representing both the coding region and the 35S promoter. These dsRNAs are substrates for production of siRNAs, which then trigger RNAi of endogenous nitrate reductase genes, TGS of other transgenes regulated by the 35S promoter, and multiple epigenetic states associated with differences in DNA methylation. Similarly, Fojtova et al. (2003) have shown that on prolonged culture of tobacco cells a neomycin phosphotransferase II transgene exhibiting RNAi gradually becomes transcriptionally inactivated. Inactivation is associated with progressive methylation that starting at the 3′ end of the transcribed region and eventually includes the 35S promoter region. In contrast, we did not observe detectable transgenerational change in methylation of the transcribed region or promoter of 35S2-YFPivs. These preliminary experiments do not rule out the possibility that changes occur at other potential methylation sites in the transgene, or at sites outside of the silenced gene, as reported for paramutation [6]. Our results also do not rule out the possibility that transcriptional activity is restricted by a chromatin-based mechanism, such as the chemical modification of histones [5,11–13,48].

Considerable evidence suggests that transcription of the target transgene above a critical threshold is required for maintenance of RNAi [40,49–52]. We speculate that gradual reduction of transcription to near this threshold results in a shift from the robust, somatically stable RNAi of type I plants to the somatically unstable RNAi of type II plants (Fig. 6). As transcription is reduced below the threshold, RNAi is completely lost and plants express a high-YFP phenotype. This model predicts that over many generations YFP mRNA expression, even in high-expressing lineages, will gradually decline and eventually produce plants silenced exclusively at the transcriptional level.

**Fig. 6.**
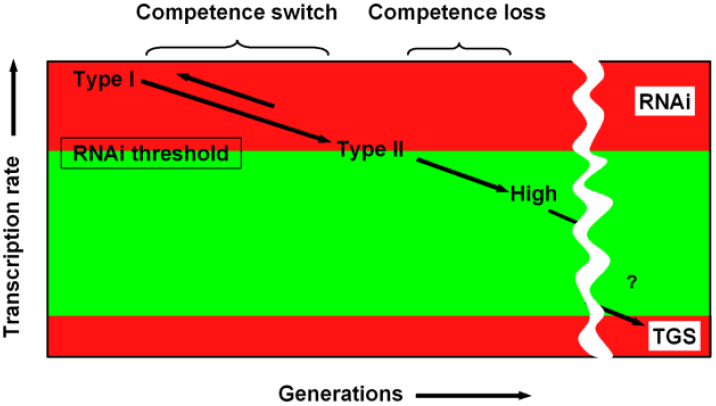
A speculative model for transgenerational effects linking RNAi and TGS. Type I plants frequently give rise to type II progeny (long arrows), and progeny that no longer show silencing (High) in successive generations. Type II plants can also give rise to revertant type I progeny at low rates (short arrow) indicating that this switch in RNAi competence is reversible and thus results from an epigenetic change. According to the model, epigenetic changes in RNAi type and loss of RNAi result from a gradual transgenerational decrease in of transgene transcription by an as yet unknown mechanism. Rates of reporter gene transcription are high in type II plants exhibiting robust RNAi. As the level of transcripts decreases to the critical threshold for maintenance of RNAi, plants exhibit unstable, type II RNAi and then the high-YFP expressing phenotype. The model predicts that eventually inactivation of transcription is complete resulting in TGS.

Although the mechanism is poorly understood, our findings emphasize the importance of transgenerational effects of RNAi in plants, which helps in interpreting these experiments and in applying RNAi to examine gene function.

## Acknowledgements

We thank Witold Filipowicz for advice and for providing Syn7; Ortrun Mittelsten Scheid for advice on nuclear run-on transcription; Barbara Hohn, Antoine Peters, Konstantina Boutsika and Claudia Kutter for critical comments; and, the Novartis Research Foundation for financial support.

## Supplementary Material

Supplementary Figure Legends

**Fig. S1.**
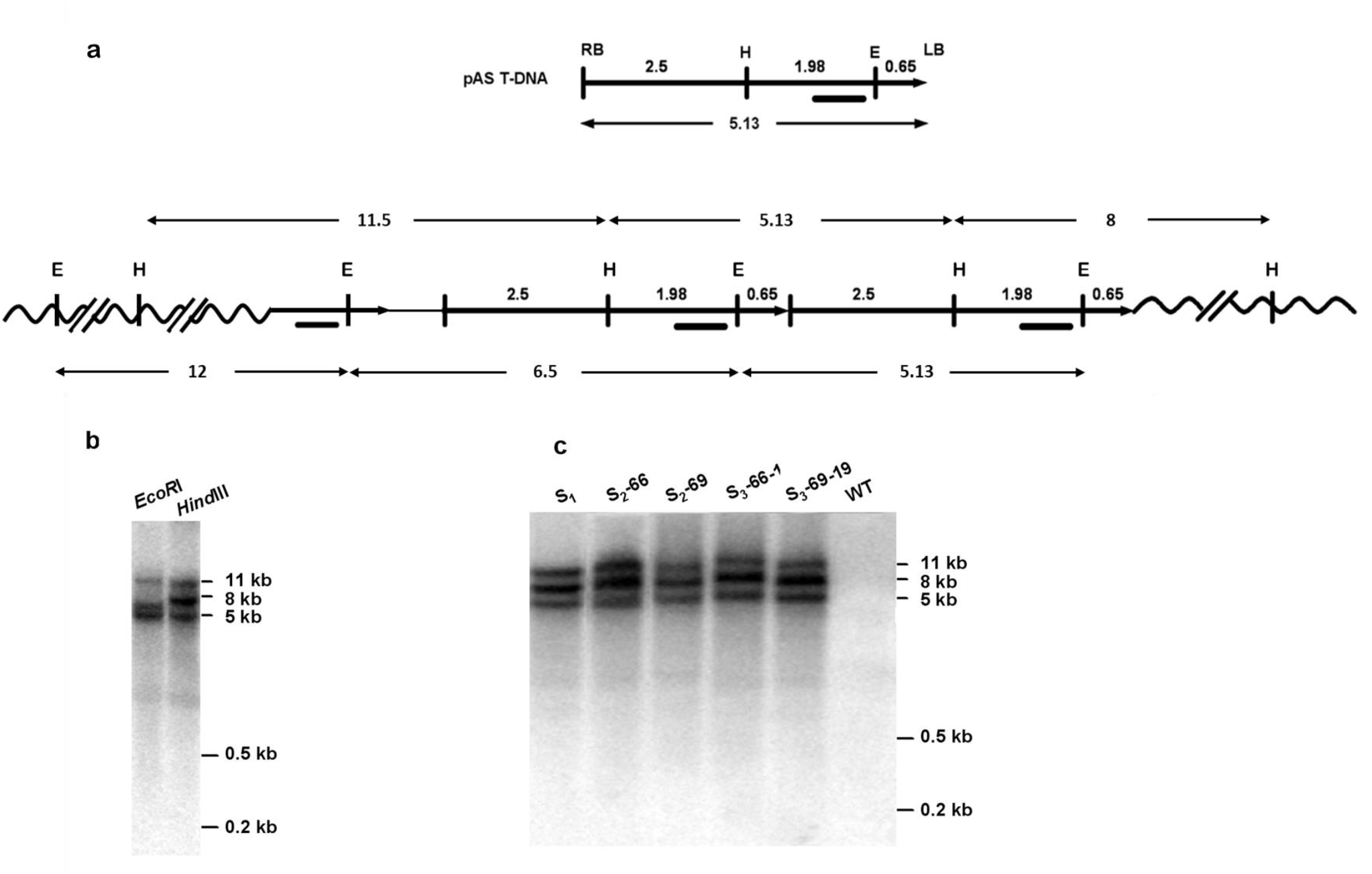
Southern blot analyses of genomic *35S_2_-YFP_ivs_* DNA. (a) A partial restriction fragment map of the pAS T-DNA and a two copy model for the T-DNA insertion in line 23. The right border (RB) and left border (LB) of the T-DNA, *Hind*III (H) and *EcoRI* (E) restriction sites, the hybridization probe used (bar), and predicted sizes of restriction fragments in kb are indicated. The position of the *Hind*III site in genomic DNA 8.87 kb from the left border was determined by Tail-PCR. (b) Southern analysis of S1 generation line 23 genomic DNA cleaved with *EcoRI* or *Hind*III. The restriction fragments found are consistent with the direct repeat model shown in (A); but, the model does not account for the additional 8 kb *Hind*III, 7 kb *EcoR*I, and 11 kb *EcoR*I fragments suggesting that a third T-DNA copy may be present at the transgene locus. (c) Southern analysis with HindIII of genomic DNA from untransformed wild-type plants (WT) and from pools of 3-5 sibling line 23 plants from the generation indicated. The pedigree of the plants used is shown in Figure 2. The very similar patterns of restriction fragments obtain with plants in successive generations indicates that the transgeneration effects observed are not due to a major change in the copy number or arrangement of the T-DNA locus.

**Fig. S2.**
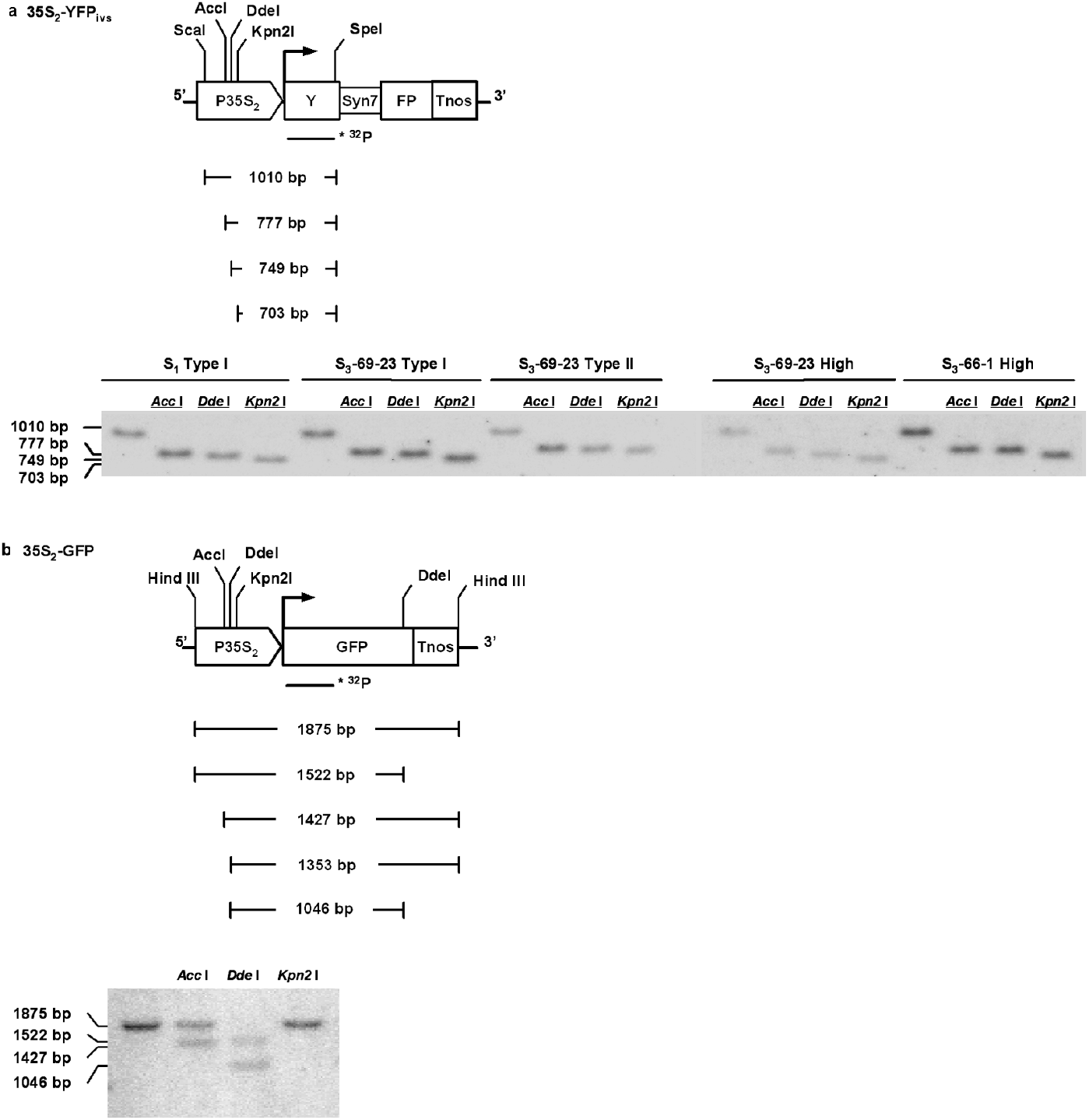
Methylation status of selected cytosines in the promoter and transcribed region of *35S_2_-YFP_ivs_* T-DNA. (a) Southern blot-analysis of genomic DNA prepared from pools of high YFP-expressing lower leaves of S_1_ type I plants as well as S_3_ generation type I, type II and high expressing plants. DNA from 5 pools of leaves was cleaved with *Scal+Spel* and in addition *Acc*I, *Ddel* or *Kpn2I* as indicated. Blots were hybridized with a ^32^P-labelled RNA probe for exon 1 of YFP_ivs_. (b) Southern blot-analysis of genomic DNA prepared from silent leaves of the 35S_2_-GFP transformant EG5, which exhibits TGS and methylation of the 35S_2_ promoter region. DNA was digested with *Hind* III and *Acc*I, *Ddel* and *Kpn2I* as indicated. Blots were hybridized with a ^32^P-labelled probe for part of the GFP coding region. Partial maps of the T-DNA constructs show the positions of relevant restriction sites, lengths of restriction fragments, and the hybridization probes used (heavy horizontal lines). Restriction enzyme cleavage was complete for all derivatives of line 23 tested indicating the absence of DNA methylation at the restriction sites tested. In contrast, DNA from line EG5 used as a positive control showed partial inhibition of cleavage by *Acc*I and *Dde*I as well as complete inhibition of cleavage by *Kpn*2I indicating the presence of partial and complete DNA methylation, respectively.

